# Dosage effect of multiple genes accounts for multisystem disorder of myotonic dystrophy type 1

**DOI:** 10.1101/658526

**Authors:** Qi Yin, Hongye Wang, Zhenfei Xie, Lifang Jin, Yifu Ding, Na Li, Yan Li, Qiong Wang, Xinyi Liu, Liuqing Xu, Kai Wang, Yanbo Cheng, Boran Chang, Cuiqing Zhong, Qian Yu, Wei Tang, Wanjin Chen, Wenjun Yang, Fan Zhang, Chen Ding, Lan Bao, Bin Zhou, Ping Hu, Jinsong Li

## Abstract

Multisystem manifestations in myotonic dystrophy type 1 (DM1) may be due to dosage reduction in multiple genes induced by aberrant expansion of CTG repeats in DMPK, including Dmpk and its neighboring genes (Six5 or Dmwd) and downstream Mbnl1. However, the direct evidences are lack. Here, we develop a new strategy to generate mice carrying multigene mutations in one step by injection of haploid embryonic stem cells with mutant Dmpk, Six5 and Mbnl1 into oocytes. The triple heterozygous mutant mice exhibit adult-onset DM1 phenotypes. With the additional mutation in Dmwd, quadruple heterozygous mutant mice recapitulate many major manifestations in congenital DM1. Moreover, muscle stem cells in both models display reduced stemness. Our results suggest that the complex symptoms of DM1 result from the reduced gene dosage of multiple genes.

## Introduction

Myotonic Dystrophy type 1 (DM1) is a complex disease with variable pathological phenotypes, disease severity, and onset ages [1-3]. The major symptoms include myotonia, muscle wasting, muscle weakness, cardiac conduction defects, cataracts and insulin resistance [3]. DM1 is a genetic disease caused by the expansion of a CTG repeat in the 3’-untranslated region of the dystrophia myotonica protein kinase (*Dmpk*) gene [4-6]. Usually, with an increasing number of the repeats, respiratory failure and mental retardation can be observed in the most severe congenital form of the disease (congenital DM1, CDM) [7-10]. A large amount of studies have shown that the triple repeat expansion not only reduces the protein or mRNA levels of *Dmpk* [11-13], but also alters the adjacent chromatin structure and reduces the expression of the neighboring genes [14-17], including downstream *Six5* [18-22] and upstream *Dmwd* [23] in DM1 cells or patients, albeit existence of controversy observations [24-26]. Moreover, numerous studies have also shown nuclear accumulation of RNA containing the expanded CUG repeats aberrantly recruits splicing regulators, such as *Mbnl1*, and forms the ribonuclear aggregates (foci) in nucleus, leading to misregulation of alternative splicing [27-32]. Mouse models carrying mutations in *Dmpk, Six5* or *Mbnl1* [33-37] could only partially recapitulate the multisystem manifestations of DM1 patients [3]. Furthermore, the defects could only be observed in homozygous mutant mice (the affected proteins were totally absent), while the affected proteins were indeed present with a decreased level in human patients. Another set of DM1 mouse models are generated by overexpression of the toxic CTG repeats [38], however, they could not fully recapitulate human phenotypes, either [38]. As an example, though the transgenic model carrying human skeletal actin (*HSA*) gene to express ∼250 untranslated CUG repeats in the muscle (*HSA*^LR^) can recapitulate multiple muscle phenotypes associated with DM1, it doesn’t exhibit other common DM1-related symptoms, such as muscle wasting and cataract [39]. Meanwhile, this model cannot mimic CDM symptoms [39]. The potential reason might be that the random insertion of transgenes in chromatin does not affect the expression of *Dmwd-Dmpk-Six5* locus, which may be also involved in the complex symptoms of DM1 [39]. Taken together, mouse DM1 models that better recapitulate the varieties of human symptoms are required to fully understand the underlying mechanism of DM1.

The multisystem symptoms of DM1 may be caused by the combination of different mechanisms, leading to down-regulation of multiple genes, including *Dmwd-Dmpk-Six5* and *Mbnl1* [40, 41]. However, the direct evidence to support the notion is missing due to the challenges to generate mice carrying multiple gene mutations simultaneously. It is time and labor consuming to generate mice carrying triple or quadruple mutations using conventional methods. Recently, mouse androgenetic haploid embryonic stem cells (AG-haESCs) have been successfully developed as sperm replacement to efficiently produce semi-cloned (SC) animals by injection into oocytes (intracytoplasmic AG-haESC injection, ICAHCI) [42-44]. AG-haESCs enable one-step efficient and stable generation of mice with multiple heterozygous mutant genes by ICAHCI of haploid cells carrying these mutations [42], allowing production of sufficient numbers of SC offspring for analyses in one generation. Thus, the haploid ESC-mediated SC technology may provide us an ideal tool to generate mouse models with multiple heterozygous mutant genes in one step to mimic the reduced expression of multiple genes in human complex diseases.

In this study, we tested our hypothesis by generating triple and quadruple heterozygous mutant mice in one step through injection of haploid ESCs (O48 cell line used in this study) carrying triple or quadruple mutant genes (*Dmpk, Six5* and *Mbnl1 or Dmpk, Six5, Mbnl1 and Dmwd*) into oocytes. Mice with triple mutations exhibited most of the major pathogenic phenotypes observed in adult-onset DM1 patients. Mice with quadruple mutations could mimic symptoms of patients from the most severe form of DM1, congenital DM1 (CDM). Interestingly, differentiation of muscle stem cells (MuSCs) is defective in both models due to stemness reduction, which recapitulates the defects in human DM1 patients and provides a novel system for screening of drugs to treat muscle problems in DM1 in the future.

## Results

### Generation of a novel DM1 model carrying mutations in *Dmpk, Six5* and *Mbnl1*

We first examined the feasibility of SC technology to generate mouse models of DM1 carrying single mutant gene by injection of haploid cells carrying mutation in *Dmpk, Six5* or *Mbnl1*, the three well-studied DM1-related genes. A haploid cell line (termed *H19*^***△*** *DMR*^*-IG*^***△*** *DMR*^-AGH or O48) that has been reported to efficiently support SC mice generation [42] was used in this study. We generated haploid ESC lines carrying mutant *Dmpk, Six5* or *Mbnl1* (referred to as ΔDmpk-O48, ΔSix5-O48 and ΔMbnl1-O48), respectively (Figure S1, S2 and S3) and found that these cells efficiently supported the generation of live SC pups carrying heterozygous mutant *Dmpk, Six5* or *Mbnl1* by ICAHCI (Table S1). Mutant SC mice grew up normally to adulthood and showed similar growth curves compared to those of wildtype (WT) SC mice produced from O48 (Figure S1, S2 and S3). Histological analysis of adult mice (4-6 month old) showed no obvious phenotypic abnormality in the muscle of *Dmpk*^+/-^, *Six5*^+/-^ or *Mbnl1*^+/-^ SC mice (Figure S1, S2 and S3).

We then set out to generate SC mice carrying triple mutations of *Dmpk, Six5* and *Mbnl1* in one step using ICAHCI. We disrupted *Six5* and *Mbnl* in ΔDmpk-O48-1 cells that have been analyzed as shown in Figure S1 and generated stable haploid cell lines carrying triple knockouts (termed ΔDSM-O48) (Figure 1a and 1b). Whole-genome sequencing analysis showed no off-target effect in ΔDSM-O48-1 cells (Table S2). ICAHCI experiments showed that ΔDSM-O48 cells (from two cell lines, i.e., ΔDSM-O48-1 and ΔDSM-O48-2) could reproducibly produce live SC pups (termed DSM-TKO SC mice) after injection into oocytes (Figure 1b and Table S1). Over 90% of DSM-TKO SC pups grew up to adulthood and showed the similar growth profiles as those of WT SC mice (Figure 1c, S4a and Table S1). We then examined muscle phenotypes commonly observed in DM1 patients, including myotonia, muscle weakness and wasting in DSM-TKO SC mice. Electromyography (EMG) tests in skeletal muscle demonstrated myotonia in adult DSM-TKO SC mice (Figure 1d). Mouse treadmill assay and grip strength test revealed that DSM-TKO SC mice displayed muscle weakness (Figure 1e and 1f); and rotarod test indicated that DMS-TKO SC mice exhibited severe motor defects (Figure 1g). Histological analysis of tibialis anterior (TA) muscles from DMS-TKO SC mice revealed several main histological hallmarks of muscles from DM1 patients, including increased number of nuclear clump and decreased fiber size (shown as the myofiber cross-sectional area (CSA)), a sign of muscle wasting (Figure 1h and 1i). Meanwhile, consistent with the clinical observations, dystrophin (Dys) expression in DMS-TKO SC mice was normal (Figure S4b). Fiber type analysis indicated an increased ratio of type I (slow) myofiber and a decreased CSA of type I in DSM-TKO SC mice (Figure S4c). Abnormalities of diaphragm muscle and small intestine were also observed (Figure S4d and S4e), implying the potential breathing and digestive dysfunctions in DMS-TKO SC mice. We next analyzed the cardiac structure and function in adult DMS-TKO SC mice because heart abnormalities are common in DM1 patients [45]. Echocardiography did not show obvious structure and functional abnormalities in 4-month-old DMS-TKO SC mice (Figure S4f and S4g). However, 12-month-old DMS-TKO SC mice exhibited significantly lower ejection fraction (Figure S4g) probably induced by increased myocardial fiber abnormalities in DMS-TKO SC mice (Figure S4h).

**Fig. 1.**
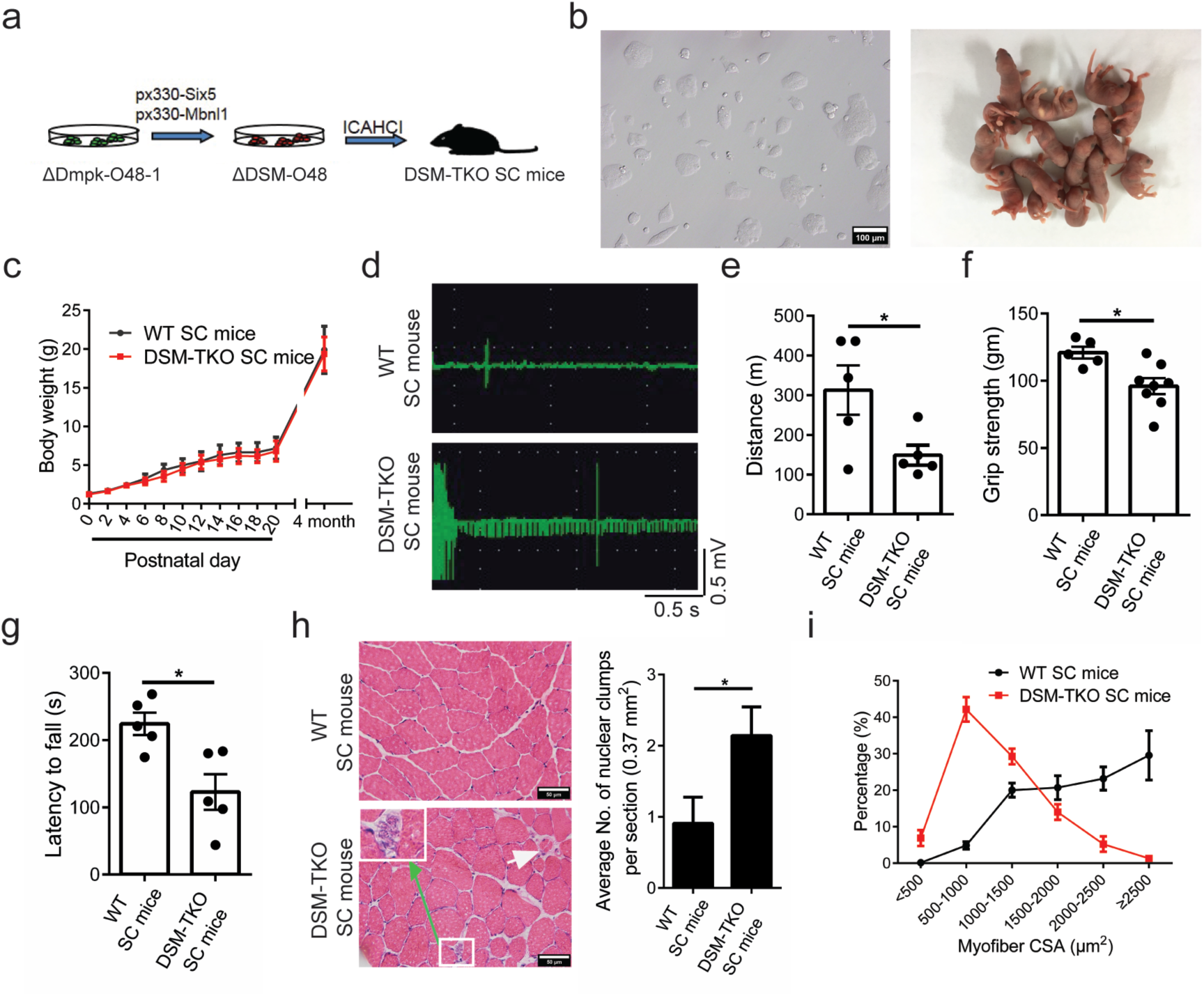
Mice carrying heterozygous mutations in *Dmpk, Six5* and *Mbnl1* exhibit typical DM1-associated muscle phenotypes. (**a)** Diagram of DSM-TKO SC mice generated by ICAHCI of ΔDSM-O48 cells. **(b)** Phase-contrast image of cultured ΔDSM-O48-1 haploid ESCs and DSM-TKO SC mice. Scale bar, 100 μm. **(c)** Body weight analysis of DSM-TKO SC mice and WT SC mice (n > 5 *per* group, mean ± s.d.). (**d)** Electromyography analysis of DSM-TKO SC mice and WT SC mice (WT SC mice, n = 3; DSM-TKO SC mice, n = 2). (**e-g)**, Muscle weakness analysis of DSM-TKO SC mice and WT SC mice was determined by Treadmill test (**e**) (n = 5 *per* group), Forelimb grip strength test (**f**) (WT SC mice, n = 5; DSM-TKO SC mice, n = 8) and Rotarod test (**g**) (n = 5 *per* group). Unpaired Student’s *t* test, \**P* < 0.05. **(h**) H&E staining of TA muscles from DSM-TKO SC mice and WT SC mice, showing nuclear clump (rectangular box) and atrophic fiber (white arrow) (WT SC mice, n = 11; DSM-TKO SC mice, n = 7). Unpaired Student’s *t* test, \**P* < 0.05. Scale bars, 50 μm. (**i)** TA muscles myofiber cross-sectional area (CSA) analysis (n = 3 *per* group).

These data demonstrate that *Dmpk*^+/-^; *Six5*^+/-^; *Mbnl1*^+/-^ SC mice can be generated in one step using haploid cells carrying triple mutations. DSM-TKO SC mice mimic the reduced dosage of three genes and display more severe pathological consequences compared to DM1 models carrying single homozygous mutant (Table S3), providing a new model of DM1. However, DSM-TKO SC mice cannot mimic several phenotypes frequently observed in DM1 patients, such as cataracts and symptoms observed in CDM patients, implying that other genes may be involved.

### *Dmwd* is involved in DM1

It has been shown that the expansion of the CTG repeats in *Dmpk* produces allele-specific effects on transcription of the two adjacent genes, *Six5* (downstream of *Dmpk*) and *Dmwd* (upstream of *Dmpk*) through changes of the local chromatin structure, leading to down-regulation of *Six5* and *Dmwd* in DM1 patients [14, 18-21, 23, 46-50]. The effects of *Six5* in DM1 have been well characterized using *Six5* knockout mice [35, 36, 51]; however, the role of *Dmwd* in DM1 has not been analyzed yet. We set out to test the function of *Dmwd* by generating haploid cells carrying mutant *Dmwd* gene using CRISPR-Cas9 (Figure 2a) [52]. Ten stable cell lines carrying mutant *Dmwd* gene (termed ΔDmwd-O48) were obtained (Figure 2b and 2c). By injecting these cells into oocytes, live SC pups could be efficiently produced (Figure 2d and Table S1). *Dmwd*^+/-^ SC mice grew up to adulthood normally (Figure 2e and Table S1). Phenotype analysis showed that the myofiber CSA was dramatically reduced in adult *Dmwd*^+/-^ mice (Figure 2f-i), suggesting that Dmwd is involved in pathological mechanism of DM1. But other symptoms of DM1, such as cardiomyocytes defects and cataracts, cannot be observed in *Dmwd*^+/-^ mice, suggesting that *Dmwd* is not sufficient to account for all the complex multisystem symptoms of DM1.

**Fig. 2.**
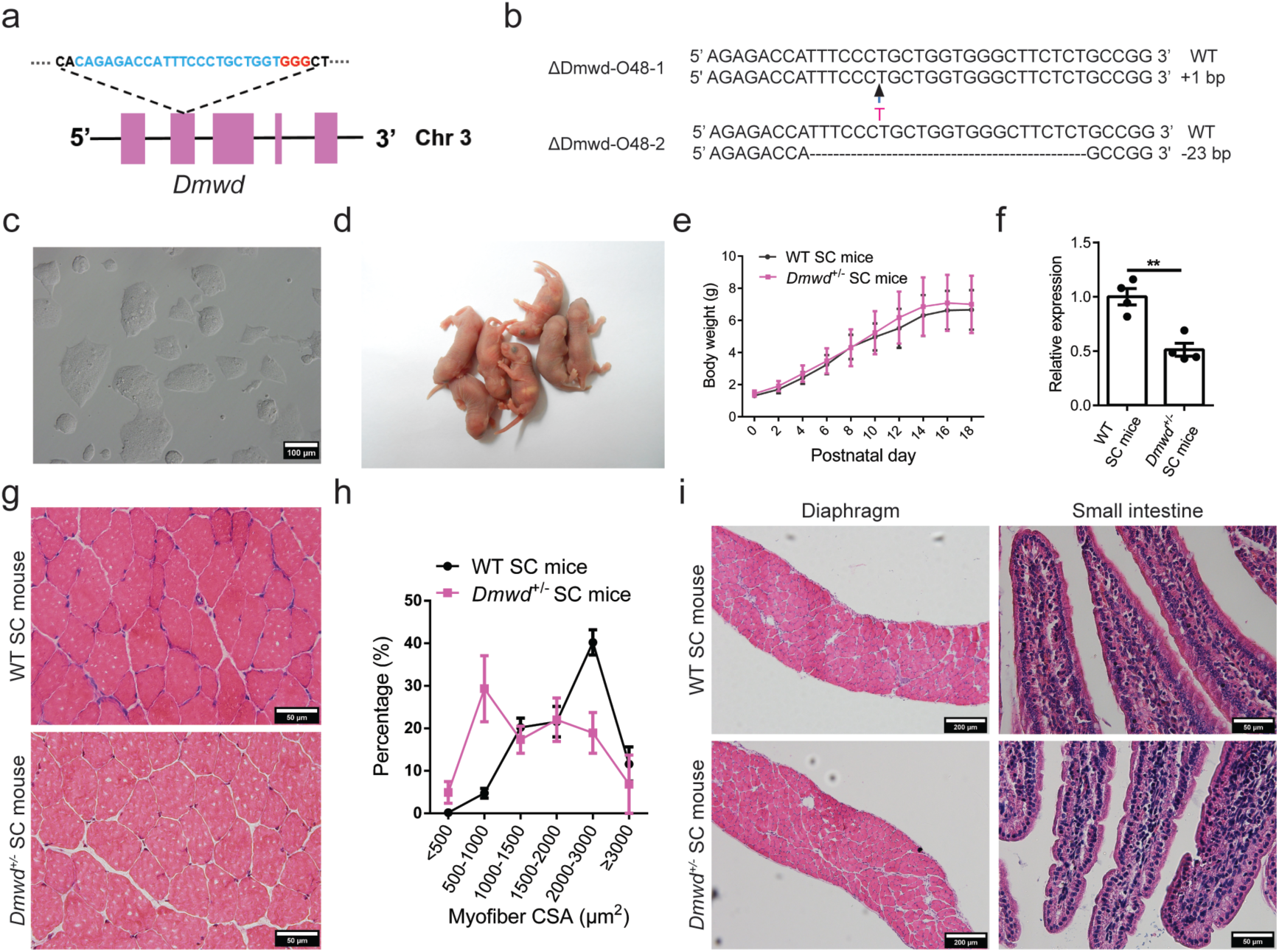
Generation of *Dmwd*^*+/-*^ SC mice through ICAHCI of haploid cells carrying mutant *Dmwd*. **(a)** Schematic of the sgRNA targeting *Dmwd*. **(b)** The sequences of the *Dmwd* gene in two cell lines (ΔDmwd-O48-1 and ΔDmwd-O48-2). **(c)** Phase-contrast image of ΔDmwd-O48-1 cell line. Scale bar, 100 μm. **(d)** Newborn *Dmwd*^*+/-*^ SC pups generated using ΔDmwd-O48-1 cell line. **(e)** Body weight analysis of *Dmwd*^*+/-*^ SC mice and WT SC mice (n > 8 *per* group, mean ± s.d.). **(f)** Transcription analysis of *Dmwd* in TA muscles showed that the transcription level of *Dmwd* was significantly reduced in *Dmwd*^*+/-*^ SC mice compared with WT SC mice (n = 4 *per* group). Unpaired Student’s *t* test, \*\**P* < 0.01. **(g)** Representative images of H&E staining of TA muscles from *Dmwd*^*+/-*^ SC mice and WT SC mice. Scale bars, 50 μm. **(h)** CSA analysis of TA muscle showed muscle wasting in *Dmwd*^*+/-*^ SC mice (n = 3 *per* group). **(i)** Representative images of H&E staining showed normal histological structure of diaphragm and small intestine in *Dmwd*^*+/-*^ SC mice Scale bars, 200 µm for diaphragm; 50 µm for small intestine.

### Generation of congenital DM1 (CDM) mice carrying quadruple mutations

Next, we tested whether the compound loss of *Dmpk, Six5, Mbnl1 and Dmwd* could give rise to a mouse model recapitulating the majority of the symptoms in DM1 patients and the more severe CDM manifestations. We further mutated *Dmwd* gene in ΔDSM-O48-1 cells and generated stable haploid cell lines carrying quadruple mutations (termed ΔDSMD-O48) (Figure 3a). Off-target analysis indicated no unexpected mutant site in tested cells (ΔDSMD-O48-2) (Table S2). ICAHCI analysis of three lines (ΔDSMD-O48-1, -2 and -3) generated *Dmpk*^+/-^; *Six5*^+/-^; *Mbnl1*^+/-^; *Dmwd*^+/-^ SC mice (DSMD-QKO SC mice) efficiently (Table S1, Figure S5a and S5b). In contrast to mice carrying the single mutation or triple mutations, around 22% of newborn DSMD-QKO SC pups with normal body weights died in a couple of hours after birth (Figure 3b). The autopsy indicated that the lung expansion failure might contribute to the neonatal death of DSMD-QKO SC pups (Figure S5c). Lung expansion failure can be caused by dysfunction of diaphragm muscle; we therefore dissected diaphragm from DSMD-QKO SC pups. The results showed obvious pathogenic alterations, including the reduced myofiber CSA and disorganized myofibers in diaphragm (Figure 3c and 3d). Moreover, immunofluorescent staining analysis showed a significant decrease of the complexity of mature neuromuscular junctions (NMJs) in DSMD-QKO diaphragm muscles (Figure 3e). Taken together, these results indicate that our DSMD-QKO SC mouse model recapitulates the postnatal respiratory insufficiency in CDM patients, which is the main cause of neonatal mortality of CDM patients [53]. They also suggest that the postnatal respiratory insufficiency in CDM patients may be due to the defects of diaphragm muscle [54].

**Fig. 3.**
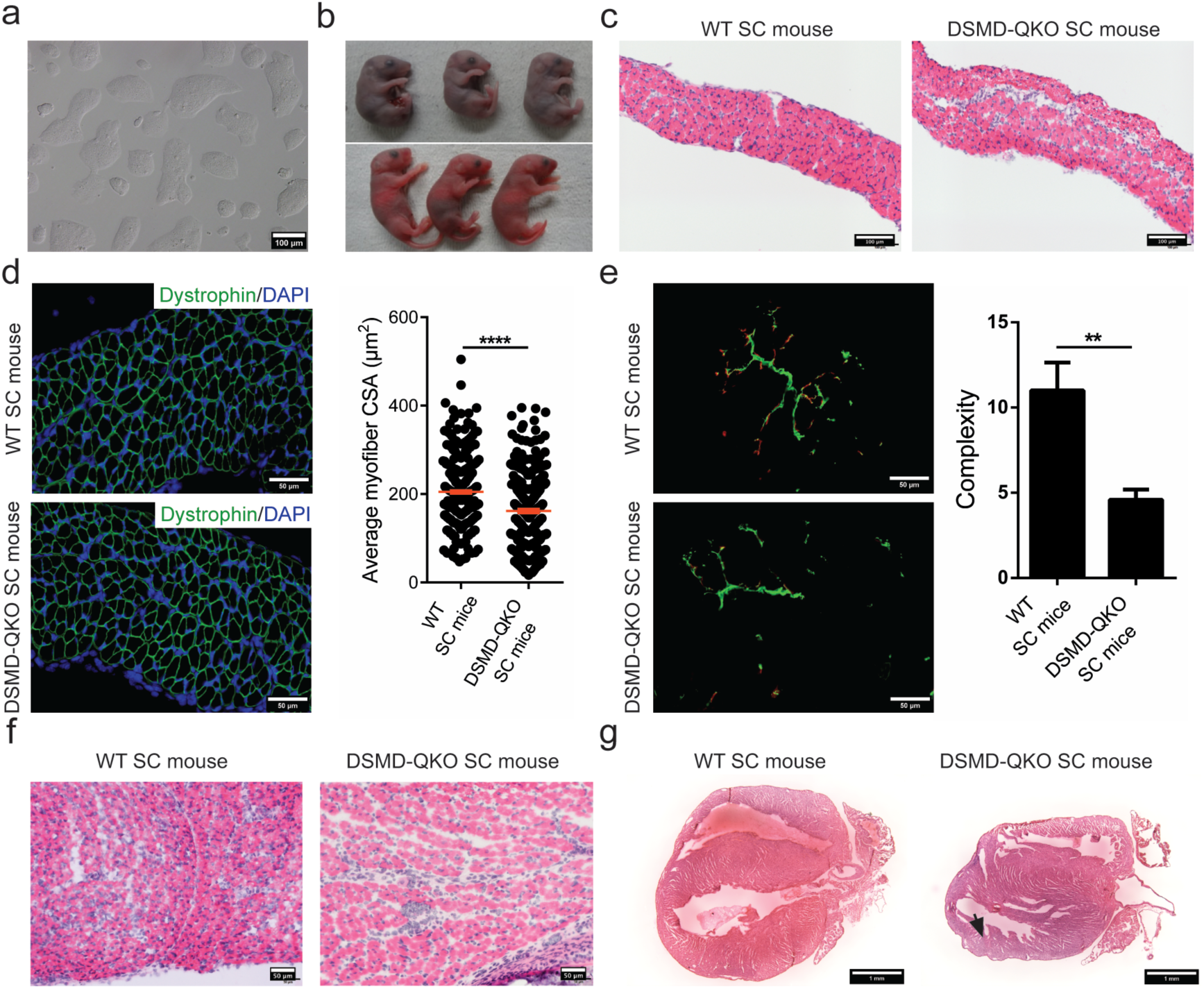
SC mice carrying heterozygous mutations in *Dmpk, Six5, Mbnl1 and Dmwd* show congenital DM1 (CDM) phenotypes. **(a)** Phase-contrast image of cultured ΔDSMD-O48-1 haploid ESCs. Scale bar, 100 μm. **(b)** 23% of newborn DSMD-QKO SC pups died during perinatal period (upper), while others survived (lower). **(c)** H&E staining of diaphragm sections from DSMD-QKO SC mice and WT SC mice (P1). Scale bars, 100 μm (**d**) Immunofluorescent staining of dystrophin and CSA analysis in diaphragm from DSMD-QKO SC mice and WT SC mice (P1) (n = 3 *per* group). Unpaired Student’s *t* test, \*\*\*\**P* < 0.0001. Scale bars, 50 μm**. (e)** Immunofluorescence staining of bungarotoxin (red) and neurofilament H (green), and complexity analysis of neuromuscular junctions of DSMD-QKO SC mice and WT SC mice (P1) (WT SC mice, n = 2; DSMD-QKO SC mice, n = 5). Unpaired Student’s *t* test, \*\**P* < 0.01. Scale bars, 50 μm. (**f**) H&E staining of TA muscle sections (P1) showing less myofiber in some DSMD-QKO SC mice (2/5). Scale bars, 50 μm. **(g)** H&E staining of heart cryo-sections (P17) of showed thinning of left ventricular posterior wall in DSMD-QKO SC mice (2/7, dark arrow). Scale bars, 1 mm.

We next characterized the growth profiles of DSMD-QKO SC mice. Interestingly, DSMD-QKO SC pups exhibited growth-retarded phenotype (Figure S5d and S5e) and 46% of them died within 3 weeks postnatally (Table S1). We sacrificed DSMD-QKO SC pups at postnatal day 2 (P2) and observed residual milk in their stomach, excluding the possibility that they died of feeding difficulties. P2 pups exhibited overt abnormal intestines (Figure S5f), consistent with the gastrointestinal abnormalities observed in CDM patients [53]. Meanwhile, some of the DSMD-QKO SC pups had abnormal talus bone development (Figure S5f), recapitulating the distinguishing feature of talipes in CDM patients [55]. Neonatal hypotonia, another typical manifestation of CDM patients [53], was also observed in DSMD-QKO SC pups (Figure S5g), probably due to less muscle fibers in the mutant pups (Figure 3f). Cardiac problem is considered to be a common cause of sudden death in patients with DM1 [45, 56]. We then performed echocardiography (ECG) test and found 2 of 7 mice (P17) exhibited ventricular premature beats (Figure S5h). Interestingly, these two pups died in a few days after ECG examination. Histological analysis of the heart showed the ventricular and atrial wall attenuation (Figure 3g). Taken together, DSMD-QKO SC mice represent a novel model that can mimic developmental defects of CDM patients, suggesting that the reduced dosage of multiple genes is responsible for the CDM phenotypes and *Dmwd* is involved in DM1 symptoms.

### Adult DSMD-QKO SC mice exhibit typical DM1 phenotypes

DSMD-QKO SC mice, once survived at weaning, could grow up to adult (Figure S5d). Western blotting analysis confirmed the reduction of protein levels of Dmpk, Six5, Mbnl1 and Dmwd in the muscle of DSMD-QKO SC mice (Figure S6a and S6b). Mass spectrometry further confirmed protein reduction in adult QKO SC mice (Figure S6c). Interestingly, adult DSMD-QKO SC mice exhibited adult DM1-associated muscle symptoms, including severe myotonia, muscle weakness and motor deficits (Figure 4a, 4b, S7a and S7b). Histological analysis indicated obvious structural abnormalities in limb muscles, including muscle wasting, and increased numbers of nuclear clumps (Figure 4c, 4d, S7c and S7d). Similar histological abnormalities could also be observed in diaphragm muscle (Figure S7e). Meanwhile, fiber type analysis showed an increased ratio and a decreased CSA of type I myofibers in DSMD-QKO SC mice (Figure 4e). Moreover, the complementary increase of type II (fast) myofiber CSA was also observed in QKO SC mice (Figure 4e), representing another severe symptom in DM1 patients [57]. In addition to muscle abnormalities, adult DSMD-QKO SC mice displayed other prominent DM1-associated features, such as distinct ocular cataracts (Figure 4f), overt small intestine abnormalities [53] and endocrine dysfunction displayed by big variation of endocrine hormone levels in different individuals, similar to the strong variations observed in patients [58] (Figure S7f-h). Although adult DSMD-QKO SC mice (4-6 month old) did not show major heart arrhythmias, they exhibited the overt functional defects and histological abnormalities in heart (Figure S7i and S7j). Interestingly, we have not observed obvious increased central nuclei in TKO and QKO mice, implying that other genes, such as MBNL2 or CUGBP1 that might be involved in DM1 patients, should be included in our mouse models in future.

**Fig. 4.**
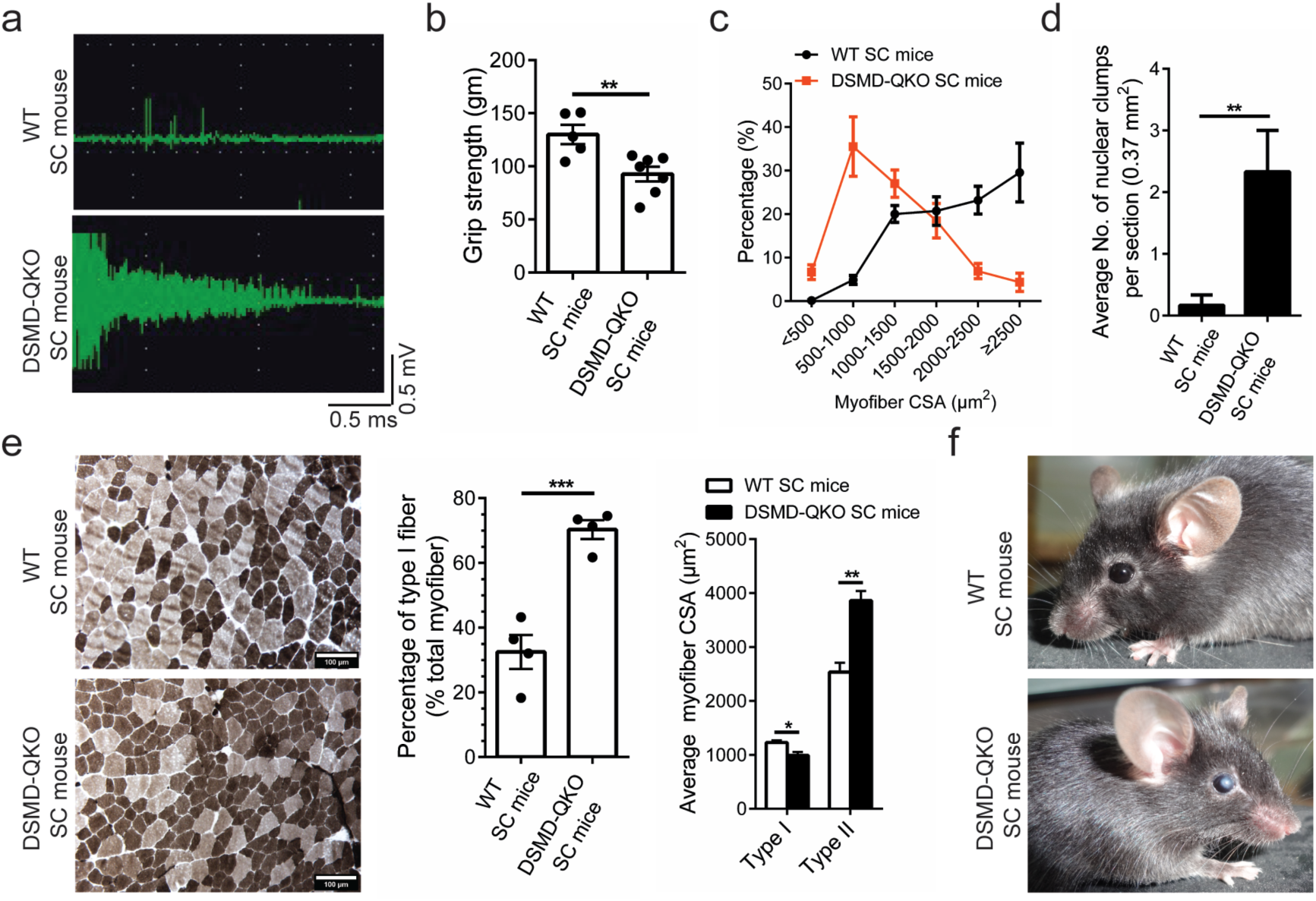
Pathogenic phenotypes in adult SC mice carrying quadruple mutations. (**a)** Electromyography analysis (WT SC mice, n = 3; DSMD-QKO SC mice, n = 5). **(b)** Forelimb grip strength test (WT SC mice, n = 5; DSMD-QKO SC mice, n = 7). Unpaired Student’s *t* test, \*\**P* < 0.01. **(c)** TA muscle CSA analysis (n = 3 per group). **(d)** Nuclear clumps analysis of TA muscle (WT SC mice, n = 6; DSMD-QKO SC mice, n = 3). Unpaired Student’s *t* test, \*\**P* < 0.01. **(e)** Representative images of ATPase staining and fiber type analysis (n = 4 *per* group). Unpaired Student’s *t*-test, \**P* < 0.05, \*\**P* < 0.01, \*\*\**P* < 0.001. Scale bars, 100 μm. **(f)** Dust-like opacities were observed in eyes of DSMD-QKO SC mice (3/7).

Since mis-splicing is a characteristic feature of DM1 and the dosage of *Mbln1* is also reduced in our models, we next analyzed splicing profiles of 7 DM1-related genes in P2, P10 and adult mice (4-month old). RT-qPCR analysis demonstrated the misregulation of *Ldb3, Serca1, m-TTN, Tmem63b, Sorbs1*, and *Spag9* mRNA splicing(Figure S8a-b), while the splicing efficiencies of *Dysf* were unchanged similar as that in the DM1 patients (Figure S8c). In addition, we also checked the splicing of *Clcn1, Ryr1, Ryr2, Tnnt2*, and *Tnnt3* mRNA, and they were showed unchanged in our mouse models (data not shown), reflecting the heterogeneity of mis-splicing in patients [59, 60] and suggesting that other genes may be involved in our models to mimic these splicing abnormalities. Taken together, DSMD-QKO SC mice can recapitulate most of the symptoms in DM1 and CDM (Table S3), providing another new model for DM1 study.

### Muscle stem cells are defective in DSM-TKO and DSMD-QKO SC mice

Previous studies have shown defective muscle stem cells (MuSCs) in DM1 patients, including aberrant stemness [61, 62], abnormal proliferation [61, 63, 64] and defective differentiation [61, 65, 66], we thus characterized the MuSCs in TKO and QKO SC mice. Immunofluorescent staining of PAX7, the MuSC marker, indicated the similar number of satellite cells in TKO, QKO (4-6 months old) and control SC mice at the same age (Figure 5a). FACS analysis further confirmed that the ratio of MuSCs (CD34+:integrin-α_7_+:CD31−:CD45−:CD11b−:Sca1−) [67] were not affected in both DM1 models (Figure S9a). We then tested the *in vitro* proliferation ability of TKO and QKO MuSCs using the long-term expansion protocol established by Fu et al [68]. The results showed that the proliferation abilities of both MuSCs were comparable to that from WT SC mice (Figure S9b).

**Fig. 5.**
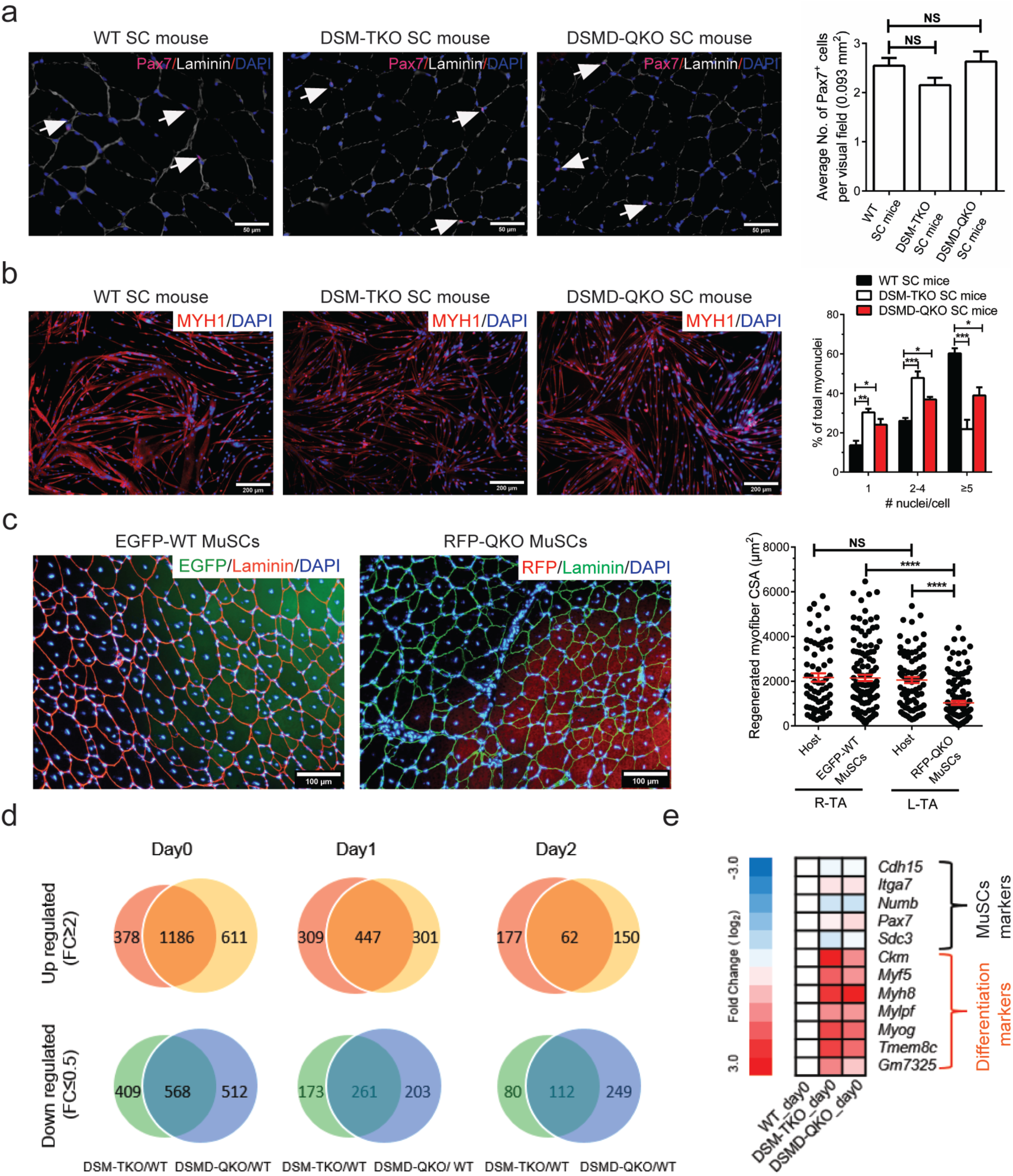
Decreased differentiation potential of MuSCs from TKO or QKO SC mice. **(a)** Representative images of Pax7 immunostaining (white arrows) of muscle sections from TKO and QKO mice (n = 3 *per* group). Scale bars, 50 µm. Unpaired Student’s *t* test. NS, no significant differences. **(b)** In vitro differentiation analysis of MuSCs (n = 3 *per* group). One-way ANOVA, \**P* < 0.05, \*\**P* < 0.01, \*\*\**P* < 0.001. Scale bars, 200 μm. **(c)** In vivo differentiation of EGFP-WT MuSCs and RFP-QKO MuSCs (n = 2 *per* group). R-TA stands for TA of right hind leg, while L-TA stands for TA of left hind leg. One-way ANOVA, \*\*\*\**P* < 0.0001. NS, no significant differences. Scale bar, 100 μm. **(d)** Venn diagrams showed that the number of common DEGs between mutant and WT samples dramatically decreased from Day 0 to Day 2. **(e)** Heat map of MuSCs markers and differentiation markers in DM1 and WT MuSCs.

We next analyzed the differentiation potentials of TKO and QKO MuSCs by culturing them in differentiation medium as reported previously [68]. The results showed that the fusion index was decreased in myofibers differentiated from both TKO and QKO MuSCs (Figure 5b). In contrast, MuSCs from a well-established transgenic mouse model expressing long repeat of CUG (*HSA*^LR^), which develops myotonia and myopathy [39], didn’t show differentiation defects *in vitro* (Figure S9c-e). To further confirm the differentiation defect was cell autonomous, we performed MuSC transplantation experiments. We generated QKO SC mice carrying RFP transgenes (termed RFP-QKO SC mice) and control SC mice carrying EGFP transgene (EGFP-WT SC mice) (Figure S10a and S10b) and then obtained MuSCs from the two strains, respectively (Figure S10c). Consistent with the above results, RFP-QKO MuSCs showed normal proliferation ability and decreased *in vitro* differentiation potential (Figure S10c). We next transplanted RFP-QKO or EGFP-WT MuSCs to TA muscles of non-fluorescent recipients. The results showed that the CSA of RFP^+^ myofibers (myofibers generated from the transplanted RFP-QKO MuSCs) was significantly smaller than the CSA of the EGFP^+^ myofibers generated from the transplanted EGFP-WT MuSCs (Figure 5c). Since both types of MuSCs shared the same recipient microenvironment, we conclude that the differentiation defects of QKO MuSCs are cell autonomous. Together, these results demonstrate decreased differentiation potential of MuSCs in TKO and QKO SC mice, mimicking another important symptom in DM1 patients.

We next attempted to reveal the underlying mechanisms by comparing the genome-wide gene expression profiles of MuSCs (Day 0) and differentiating cells (Day 1 and Day 2) between mutant and normal SC mice (Figure S11a). Clustering analysis indicated that cells with the same differentiation stage rather than with the same genotype exhibited higher correlation (Figure S11b), suggesting that TKO and QKO mutations do not dramatically change the cell fate. Interestingly, the total number of differentially expressed genes (DEGs) between mutant cells and WT cells dramatically decreased after induction of differentiation (Figure 5d), implying that the major differences existed at the stem cell stage. Enrichment analysis of DEGs on Day 0 showed that muscle differentiation-related genes (such as muscle development and contraction) were up-regulated in mutant MuSCs compared to WT cells (Figure 5e and S11c-S11e). In contrast, the expression levels of stem cell markers were similar (Figure 5e and S11d-S11e). Nevertheless, qRT-PCR analyses indicated that the expression levels of the muscle-differentiation-related genes were comparable in differentiating cells from mutant and wild-type MuSCs (Figure S11f). These results demonstrate that MuSCs of TKO and QKO SC mice are at a more committed state [69]. The premature expression of differentiation genes at stem cell stage suggests the partial loss of stemness and may account for the decreased differentiation potential of mutant MuSCs *in vitro* and *in vivo*.

## Discussion

Mutations or expression changes occur in multiple genes in most of the diseases, especially in those multisystem syndromes. Generation of mouse models carrying multigene mutations to faithfully recapitulate the complex symptoms is the key to elucidate the mechanism of these syndromes. However, it is time consuming and labor intensive to do so using the conventional methods, leading to the lack of proper animal models for many syndromes. In this proof-of-concept study, we generated mouse models to mimic the reduced expression of multiple genes in human DM1 in one step by ICAHCI of haploid ESCs carrying three or four mutant genes. Interestingly, mice carrying heterozygous mutant *Dmpk, Six5* and *Mbnl1* (DSM-TKO SC mice) exhibit typical symptoms of adult-onset DM1. With the additional haploid mutation in *Dmwd*, DSMD-QKO SC mice faithfully recapitulate the pathogenic phenotypes of the more severe CDM patients. These results suggest that dosage effects of multiple genes affected by different mechanisms, including RNA toxicity and local chromatin changes, are the major pathogenic factors in DM1 and the severity of the disease depends on the number of genes affected. In patients, longer CTG repeats are associated with more severe symptoms and earlier disease onset; however, the underlying mechanism is unknown. Our results may provide potential explanation: besides abnormal-RNA-induced down-regulation of *Mbnl1* as suggested by the RNA disease model, local chromatin structure changes induced by the aberrant expansion of CTG repeats in 3’-untranslated region of *Dmpk* gene result in decreased expression of its downstream *Six5* (DSM-TKO SC mice); with an increasing number of repeats, the expression of its upstream *Dmwd* is also affected (mimicked by DSMD-QKO SC mice), leading to more severe and early onset DM1 (Table S3). The longer the CTG repeat expansion is, the expression of more genes may be affected. Interestingly, MuSC stemness is affected in our DM1 mouse models, consistent with the observations in both DM1 and CDM patients [61, 63, 70] and recent observations in homozygous mice carryin *Mbnl3* gene depletion [26]. Taken together, our study provides the direct evidence to support the hypothesis that expansion of the CTG repeats affects the expression level of multiple genes, leading to complex pathogenic phenotypes in DM1 patients. Meanwhile, TKO and QKO mouse models may provide a novel platform for drug screening to treat DM1.

Our method paves the road towards generating a series of mouse models carrying multiple gene mutations in a short time to mimic different stages and severity of the complex syndromes. A potential application of the approach is to produce mice carrying multiple modifications in candidate loci that have been identified in high-throughput studies or genetic screenings to mimic clinical manifestations of multigenic diseases. In summary, we have established novel DM1 mouse models by one-step injection of haploid cells carrying three or four mutant genes into oocytes, providing suitable models to investigate the molecular mechanisms underlying complex manifestations and perform drug screening. Future analysis of the TKO and OKO mice will reveal more genes involved in the complex manifestations of DM1. Meanwhile, we hope that our haploid ESC-mediated SC technology will promote modeling of other complex diseases in mice.

## Acknowledgements

*HSA*^*LR*^ mice were obtained from Dr. Charles Thornton in the Wellstone Center at the University of Rochester (NIH NS048843). We thank Y. Chen and X. Ding from the Core Facility for Stem Cell Research at SIBCB for support with cell cultures and M. Chen from Core Facility for Chemical Biology for support with small-molecule screen. This study was supported by Genome Tagging Project and grants from the Ministry of Science and Technology of China (2017YFA0102700, and 2014CB964700), the National Natural Science Foundation of China (31530048, 31601163, 81672117, 91649104, and 31671536), Chinese Academy of Sciences (XDB19010204, OYZDJ-SSW-SMC023, and KJZD-EW-L13), CAS-CSIRO cooperative Research Program (GJHZ1504), Shanghai Municipal Commission for Science and Technology (16JC1420500, 17JC1400900 and 17411954900).

## Author Contributions

J.L., P.H. and Q.Y. conceived the projects. Q.Y., Z.X. and C.Z. contributed to the haploid cell generation. Q.Y. and Z.X. performed ICAHCI. Q.Y., H.W., L.J., Y.D. and N.L. characterized the muscle phenotypes. Q.Y., Y.L. and B.Z. analyzed heart phenotypes. Q.W and L.B. performed rotarod experiments. Q.Y., Z.X., Q.Y., Y.C. B.C.,and W.T. performed embryo transplantation. X.L., L.X. and W.C. performed EMG. K.W. analyzed RNA-seq data. J.L., P.H., Q.Y., and D.L. wrote and revised the manuscript.

## Materials and Methods

### Experimental mice

All animal procedures were carried out in accordance with the guidelines of the Shanghai Institute of Biochemistry and Cell Biology (SIBCB). All mice were housed in specific pathogen-free facilities of SIBCB. Oocytes for micromanipulation were obtained from female mice of B6D2F1 (C57BL/6 × DBA/2) background. *HSA*^*LR*^ mice (FVB/n background) and *IL2r*^−*/*−^ mice (C57BL/6 background) were purchased from the Jackson Laboratory.

### Derivation of gene-modified haploid cell lines

CRISPR-Cas9-mediated Gene manipulation was performed as previously described methods [52, 71]. Briefly, oligoes for different genes (*Dmpk, Six5, Mbnl1* and *Dmwd*) were synthesized and ligated in px330-mcherry plasmid. Constructed plasmids were then transfected into haploid cells (O48), which were cultured in the ESC medium plus 3 μM CHIR99021, and 1 μM PD0325901 (ES+2i). The haploid cells were enriched through FACS and plated in one well of the 6-well plate at low density to obtain single-cell clone 24 h after transfection. 6-7 days after plating, the single-cell clones were picked and separated into two parts, one for passaging and the other for sequencing to determine the genotype. For generation of EGFP-O48 and RFP-ΔDSMD-O48 haploid ESCs, the haploid ESCs were transfected with PiggBac plasmids (CAG-EGFP or CAG-RFP).

### ICAHCI

To produce semi-cloned (SC) embryos, the haploid ESCs arrested in M stage by cultured in medium containing 0.05 μg/ml demecolcine were trypsinized and resuspended in hepes-CZB medium, then injected into oocytes using a Piezo-drill micromanipulator (prime technology Ltd) as described previously [72]. The SC embryos were cultured in KSOM medium with amino acids at 37°C under 5% CO_2_ in air. The 2-cell stage embryos were transferred into oviduct of pseudopregnant ICR females at 0.5 dpc with 15–20 embryos per side. Recipient mothers naturally delivered the SC pups.

### Immunostaining

Dissected tissues from diaphragm, heart and tibialis anterior (TA) muscle were put into the OCT (Leica) directly, and then frozen in liquid nitrogen for 15 s. The frozen tissues could be stored in -80°C refrigerators until sectioning. For whole heart longitudinal sections, the fresh dissected tissues were washed with PBS and fixed with 4% paraformaldehyde (PFA) at 4°C for 30 min. After dehydrated with 30% sucrose solution, the samples were embedded in OCT. The protocol for Pax7 (DHSB, 1:200) staining was carried out according to previously described protocols [68, 73]. The immunostaining for dystrophin was perform using anti-dystrophin (abcam, ab15277) as primary antibody (1:400) and Alexa 488-conjunated anti-mouse antibodies (Invitrogen) as secondary antibody. All images were acquired on Leica SP8 confocal microscope. For NMJs structural analysis, the frozen transverse sections of diaphragm were stained with a-bungarotoxin conjugate with Alexa Fluor 594 (Invitrogen, B13423) for acetylcholine receptors, and then incubated with chicken ployclonal antibody for neurofilament H (EnCor Biotechnology, CPCA-NF-H, 1:2000) and followed by incubation with Alexa 488 labeled secondary antibody.

### Histology

Frozen transverse sections (10 μm) of TA and diaphragm muscles were stained with hematoxylin and eosin (H&E). Fiber cross-sectional areas were measured using ImageJ. ATPase staining was performed as described previously [74]. Briefly, transverse sections were incubated in pH 4.6 solution for 5-15 min, followed by staining with ATP solution. After washed with medium containing 1% w/v CaCl_2_, 2% w/v CoCl_2_, 1:20 solution of 0.1 M Sodium Barbital and 1 % v/v solution of ammonium sulfide, the sections were dehydrated within ascending alcohols. CSA of Type I and type II fiber were measured using ImageJ. For small intestine, freshly dissected tissues were flushed with PBS and 4% PFA, followed by fixation in 4% PFA overnight at 4°C, dehydrated, embedded in paraffin, and cut at 5 μm thickness.

### Grip strength, treadmill, rotarod test and righting assay

The forelimb grip strength of mice (4-month-old) was assessed using a grip strength meter. Every mouse was tested for five times and the arithmetic average value of the records shall be as the measured value. For treadmill test, following familiarization test (6 m/min for 3 min), the mouse (4-month-old) was placed on a treadmill with a start speed of 10 m/min. Running speed was increased by 2 m/min every two minutes until 30 m/min, and the distance was recorded when the mouse could no longer run. For rotarod performance test, the mouse (4-month-old) was placed on a rotating rod at uniform motion (10 rpm) for 5 min and then an accelerating rotarod, which started with 4 rpm and gradually increased to 40 rpm in 5 minutes. The mice were pre-trained for 2 days for adaptation. The duration time on the rotarod before the mice fell off was recorded. As for righting assay, the mice at P5 were placed on its back to determine if the mice could right themselves to normal posture with four paws on the ground.

### MuSC isolation, expansion and *in vitro* differentiation

MuSCs were isolated according to previously described methods [67, 68, 75]. MuSCs were cultured on collagen-coated dish in T cell conditional medium for long-term expansion according to our reported protocol [68]. MuSCs were differentiated in differentiation medium with 2% horse serum (Sigma). After differentiated for 48 h, the cells were stained with anti-Myh antibody (Merck Millipore, 05-716), and counterstained with DAPI for counting the number of the nuclei in one fiber.

### MuSC transplantation (*in vivo* differentiation)

MuSC transplantation was performed according to described protocols [76, 77] with some modifications. Briefly, both hindlimbs of host *IL2r*^−*/*−^ mice were exposed to 18 Gy gamma radiations. Twenty-four hours before MuSC transplantation, 10 μM CTX (15 μl) is intramuscularly injected into mouse TA muscle via a 29 G insulin syringe. 5 × 10^5^ MuSCs were resuspended with 30 μl of PBS and injected into TA muscle using 29 G insulin syringe. TA muscles were collected in 4 weeks after cell transplantation and used for histological analysis.

### EdU labeling

MuSCs were cultured in MuSC medium with EdU solution (Invitrogen, final concentration, 10 μM) for 2 h at 37 °C, followed by fixation with 3.7% PFA, permealized with 0.5% Triton X-100, reacted with Click-iT reaction cocktail and counterstained with Hoechst 33342.

### Skeletal preparation and staining

Skeletal preparations were stained with alcian blue and ARS. Briefly, the mice were executed, eviscerated and skinned. The samples were dehydrated with 95% ethanol for 3 days, and then were stained in Alcian blue solution for 3 days. After fixed and cleared with 95% ethanol for three times (1.5 h for each), the samples were treated with 2% KOH for 3–4 h and stained with ARS solution for another 3–4 h. After staining, skeletons were cleared in 1% KOH/20% glycerol and stored in 100% glycerol up to several years.

### ELISA

Insulin, PTH and TSH levels in blood were measured by ELISA kit (Ebioscience) according to manufacturer’s instructions. The whole blood was collected from Retro-orbital Bleeding.

### Electromyography (EMG) and echocardiography

EMG was performed under halothane anesthesia using 30-gauge concentric needle electrodes, with sampling of distal muscle group in left hindlimb. For echocardiography analysis, after removing the hair from chest using depilatory cream, the mouse was anesthetized and fixed on Vevo2100 (VisualSonics) with tighting the limbs on the mental detector to detect electrocardiography (ECG). The echocardiography was recorded according to manufacture’s instructions.

### Western blotting

Dissected tissues were frozen with liquid nitrogen and stored in -80°C refrigerators until protein extraction. The homogenization of the tissue was done by vortex in cell lysis buffer. After separated on SDS-PAGE gels, the lysate was transferred to PVDF membranes and incubated with primary antibodies (anti-Mbnl1, abcam, ab108519; anti-SIX5, abcam, ab113064; anti-DMWD, santa cruz, sc-167638; anti-DMPK, santa cruz, sc-13612) at 4°C overnight, followed by incubation with secondary antibodies.

### RNA-seq and gene expression analysis

MuSCs (Day 0) and differentiating cells (Day 1 an Day 2) were collected and referred to RNA extraction using TRIZOL Reagent (Invitrogen). The total RNA was processed for RNA-seq by TruSeq RNA sample pre kit (Illumina) according to the manufacturer’s instructions, and checked for a RIN number (≥ 7) to inspect RNA integrity by an Agilent Bioanalyzer 2100 (Agilent technologies). Qualified RNA was further purified by RNAClean XP Kit (Beckman Coulter, Inc.) and RNase-Free DNase Set (QIAGEN). Adaptors as well as low quality base pairs were trimmed. The sequencing was performed using Illumina Hiseq X TEN and then the preprocessed reads were aligned to the protein-encoding gene of mouse reference genome (NCBI Mus musculus assembly GRCm38.p5) using hisat2 (version 2.0.5). Transcriptome profiling was determined in units of RPKM (Reads Per Kilobase per Million mapped reads) using locally developed Perl script.

### Quantitative RT-PCR analysis

Total RNA was extracted from dissected tissues according to the manufacturer’s instructions. 0.5 μg of total RNA was reverse transcribed using a First Strand cDNA Synthesis kit (TOYOBO). qPCR was carried out with THUNDERBIRD SYBR qPCR Mix (Toyobo) on a CFX connect Real-Time System (BIO-RAD). Q-PCR primers were listed in Table S3. For alternative splicing analysis, the primers, which could separate the splice alternatives of target genes [78], were used to amplify the RNA variants, followed by running on 2% agarose gel or SDS-PAGE to calculate the splicing efficiency (percent of spliced in).

### Statistical analysis

Quantitative values are presented mean ± s.e.m, unless noted otherwise. Statistical differences between groups were determined by using GraphPad Prism 6 using unpaired two-tailed *t*-test. No statistical method was used to predetermine sample size. Investigators were not blinded to outcome assessment.

### Accession number

All RNA-seq data sets are available through GEO under the accession number GEO: GSE103841.

