## Supplementary material for "Dosage effect of multiple genes accounts for multisystem disorder of myotonic dystrophy type 1": Supplymentary Data

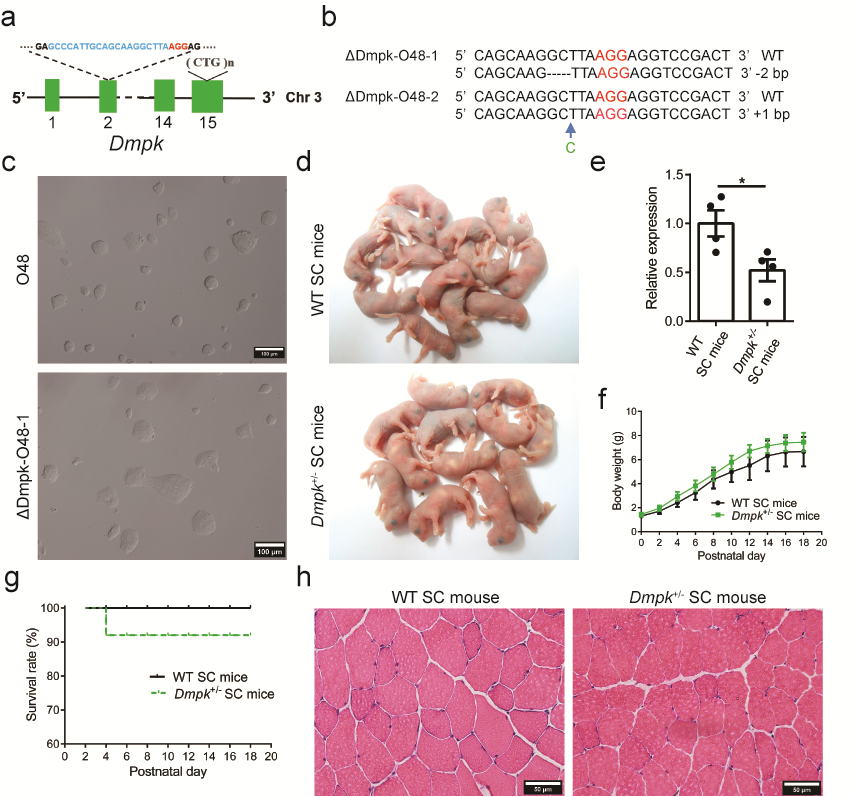

**Fig. S1** Generation of *Dmpk^+/-^* SC mice through ICAHCI of haESCs carrying mutant *Dmpk*. **(a)** Schematic of the sgRNA targeting *Dmpk*. **(b)** The sequences of the *Dmpk* in two cell lines (ΔDmpk-O48-1 and ΔDmpk-O48-2). **(c)** Phase-contrast image of ΔDmpk-O48-1 and O48 cell lines. Scale bars, 100 μm. **(d)** Newborn SC pups from ΔDmpk-O48-1 and O48 cell lines. **(e)** Transcription analysis of *Dmpk* in *Dmpk^+/-^* SC mice and WT mice (n = 4 *per* group) showing the expression level of *Dmpk* was significantly reduced in *Dmpk^+/-^* mice compared with WT SC mice. Unpaired Student’s *t* test, **P* < 0.05. **(f)** Body weight analysis of *Dmpk*^+/-^ SC mice and WT SC mice (n > 10 *per* group, mean ± s.d.). **(g)** Survival curve of *Dmpk^+/-^* SC mice and WT SC mice (n > 10 *per* group). **(h)** Representative images of H&E staining of TA muscles from *Dmpk^+/-^* SC mice and WT SC mice. Scale bars, 50 μm.

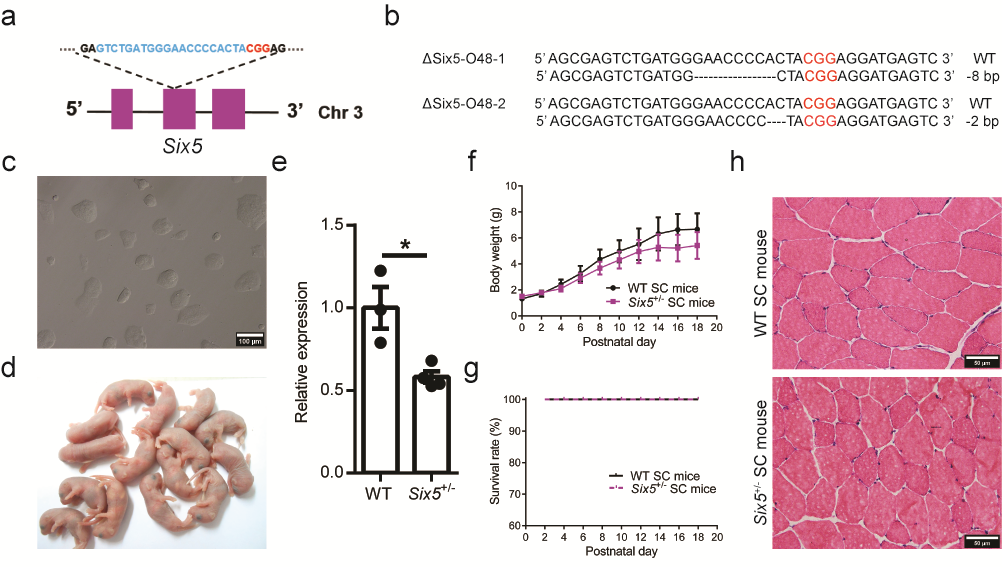

**Fig. S2** Generation of *Six5^+/-^* SC mice through ICAHCI of haploid ESCs carrying mutant *Six5*. **(a)** Schematic of the sgRNA targeting *Six5*. **(b)** The sequences of the *Six5* gene in two cell lines (ΔSix5-O48-1 and ΔSix5-O48-2). **(c)** Phase-contrast image of ΔSix5-O48-1 cell line. Scale bar, 100 μm. **(d)** Newborn SC pups generated from ΔSix5-O48-1 cell line. **(e)** Transcription analysis of *Six5* in *Six5^+/-^* and WT SC mice (*Six5^+/-^* SC mice, n = 4; WT SC mice, n = 3) showing the expression level of *Six5* was significantly reduced in *Six5^+/-^* SC mice compared with WT SC mice. Unpaired Student’s *t* test, **P* < 0.05. **(f)** Body weight analysis of *Six5^+/-^* SC mice and WT SC mice (n > 4 *per* group, mean ± s.d.). **(g)** Survival curve of *Six5^+/-^* SC mice and WT SC mice (n > 4 *per* group). **(h)** Representative images of H&E staining of TA muscles from *Six5^+/-^* SC mice and WT SC mice. Scale bars, 50 μm.

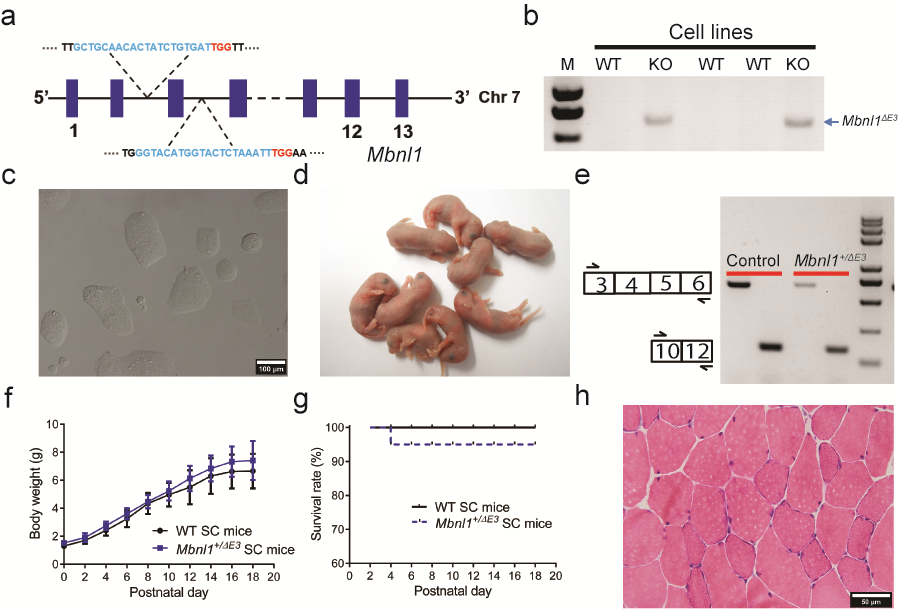

**Fig S3** Generation of *Mbnl1^+/ΔE3^* SC mice through ICAHCI of haploid cells carrying mutant *Mbnl1*. **(a)** Schematic of two sgRNAs for removing exon 3 of *Mbnl1*. **(b)** Genotyping analysis of *Mbnl1*^ΔE3^-O48 cell lines. **(c)** Phase-contrast image of *Mbnl1^ΔE3^*-O48-2 cell line. Scale bar, 100 μm. **(d)** Newborn SC pups generated from Mbnl1^ΔE3^-O48-2 cell line. **(e)** RT-PCR of *Mbnl1* in TA muscles from *Mbnl1^+/ΔE3^* SC mice and WT SC mice. **(f)** Body weight analysis of *Mbnl1^+/ΔE3^* SC mice and WT SC mice (n > 8 *per* group, mean ± s.d.). **(g)** Survival curve of *Mbnl1^+/ΔE3^* SC mice and WT SC mice (n > 8 per group). **(h)** Representative image of H&E staining of TA muscles from *Mbnl1^+/ΔE3^* SC mice. Scale bar, 50 μm.

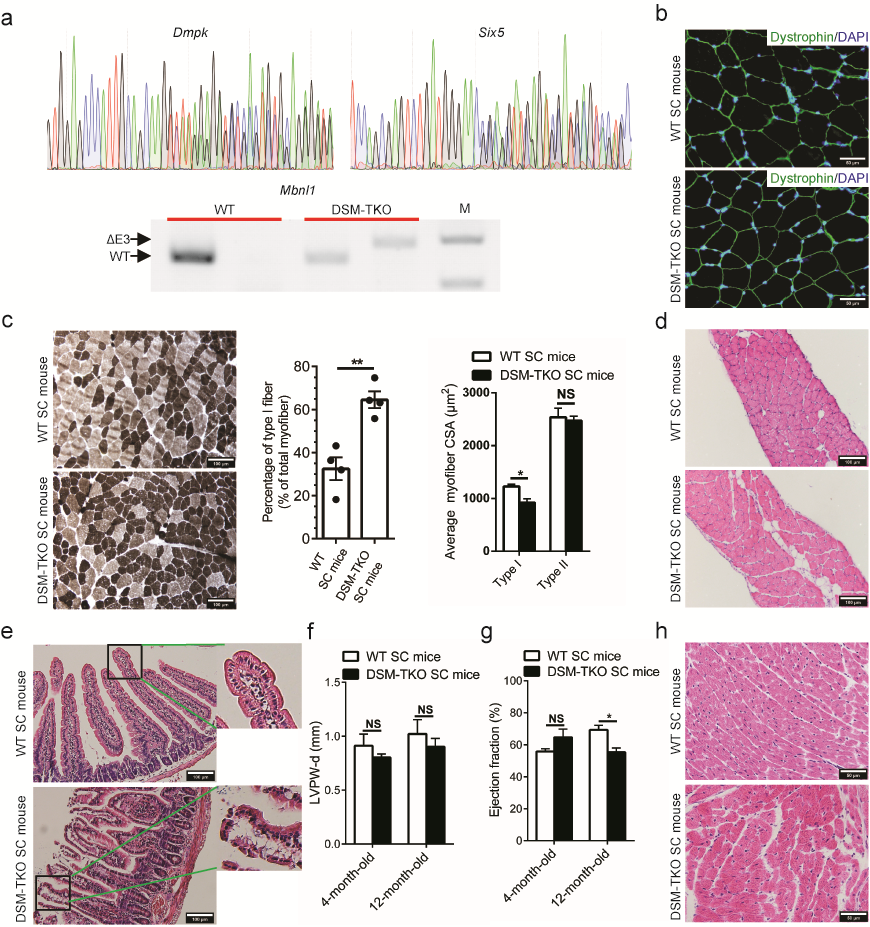

**Fig. S4** DM1-associated phenotypes in DSM-TKO SC mice. **(a)** The genotyping of the DSM-TKO SC mice. **(b)** Immunofluorescent staining indicated equivalent expression of dystrophin in DSM-TKO SC mice and WT SC mice. Scale bars, 50 μm. **(c)** ATPase staining analysis indicated increased ratio of typeⅠ(black) myofiber and reduced average CSA of typeⅠfiber in DSM-TKO SC mice (n = 4 *per* group). Unpaired Student’s *t* test, **P* < 0.05, ***P* < 0.01. NS, no significant differences. Scale bars, 100 μm. **(d)** Representative images of H&E staining of diaphragm showed increased fat cell invasion in DSM-TKO SC mice. Scale bars, 100 μm. **(e)** Representative images of H&E staining of small intestinal sections. Boxes in left panels are magnified in right panels, showed disorganization of intestine villi in DSM-TKO SC mice. Scale bars, 100 μm. **(f)** IVPW-d thickness (diastole) of DSM-TKO SC mice and WT SC mice at different ages (n = 5 *per* group). IVPW: left ventricular posterior wall. Unpaired Student’s *t* test. NS, no significant differences. **(g)** Reduced ejection fraction was observed in 12-month-old DSM-TKO SC mice (n = 5 *per* group). Unpaired Student’s *t* test, **P* < 0.05. NS, no significant differences. **(h)** Representative images of H&E staining of cardiac muscle showed myocardial disarray in 12-month-old TKO SC mice. Scale bars, 50 μm.

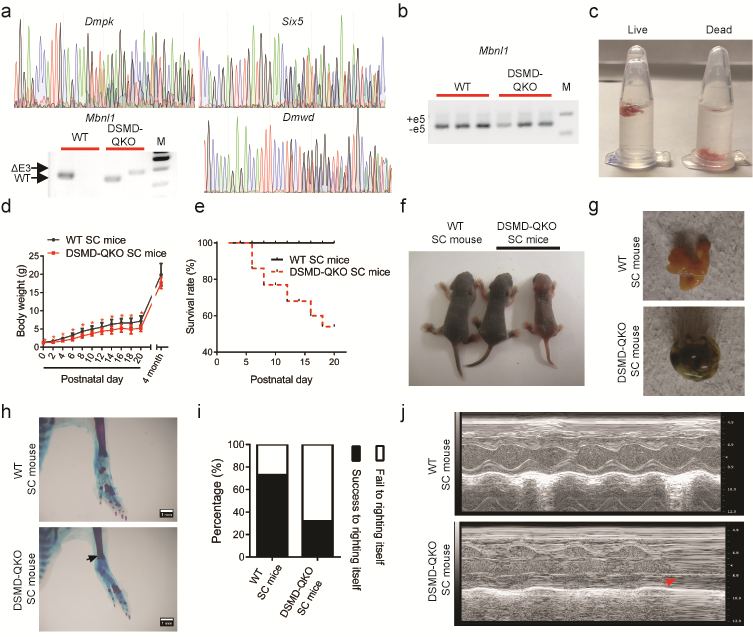

**Fig S5** CDM1-related phenotypes DSMD-QKO SC mice before weaning. **(a)** The genotyping of the DSMD-QKO SC mice. **(b)** Alterative splicing analysis of *Mbnl1* exon5 in DSMD-QKO SC mice showing normal pattern compared with WT SC mice. **(c)**The lung dissected from dead and live DSMD-QKO SC pups were dropped into water (n = 3 per group). The lungs of dead DMSD-QKO SC pups never inflated with air sinks to the bottom. **(d)** Body weight analysis of DSMD-QKO SC mice and control SC mice revealed delayed growth in DSMD-QKO SC mice (n > 10 *per* group). Unpaired Student’s *t* test, **P* < 0.05. **(e)** Survival curve of DSMD-QKO and WT SC mice (n > 10 *per* group). **(f)** Developmental failure of intestines (3/16) were observed in DSMD-QKO SC pups. **(g)** Smaller talus bone (3/7, black arrows) in DSMD-QKO SC pups compared with WT SC mice. Scales bars, 1 mm. **(h)** Representative image of DSMD-QKO and WT SC pups at postnatal day 5 (P5). The right one is a DSMD-QKO pup with growth-retarded phenotype. The middle one is a DSMD-QKO pup with normal growth. The left one is a control SC pup. **(i)** Righting assay showed that more DSMD-QKO SC mice (P5) failed to right themselves than control SC mice (WT SC mice, n = 15; DSMD-QKO SC mice, n = 22). **(j)** Echocardiography experiments showed ventricular premature beats in DSMD-QKO (2/7). Red arrow indicates ventricular premature beats.

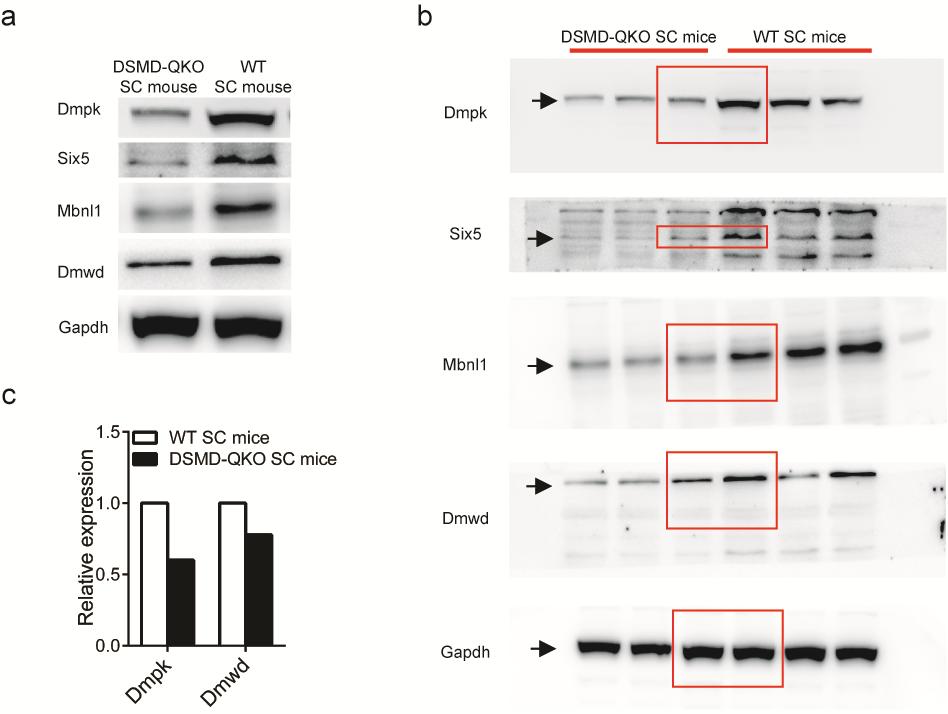

**Fig. S6** Proteins expression analysis of *Dmpk*, *Six5*, *Mbnl1* and *Dmwd* in the tissues of adult DSMD-QKO SC mice and WT SC mice. **(a)** Representative image of western blotting of Dmpk, Six5, Mbnl1, Dmwd and Gapdh in TA muscles from adult DSMD-QKO SC mice and WT SC mice. **(b)** The original gel of (a). The arrowheads indicated the target protein and the red box indicated the cutting position of (a). **(c)** Semi-quantitative analysis of the expression of *Dmpk* (tissue: stomach) and *Dmwd* (tissue: brain) of DSMD-QKO SC mice and WT SC mice using mass spectrometry.

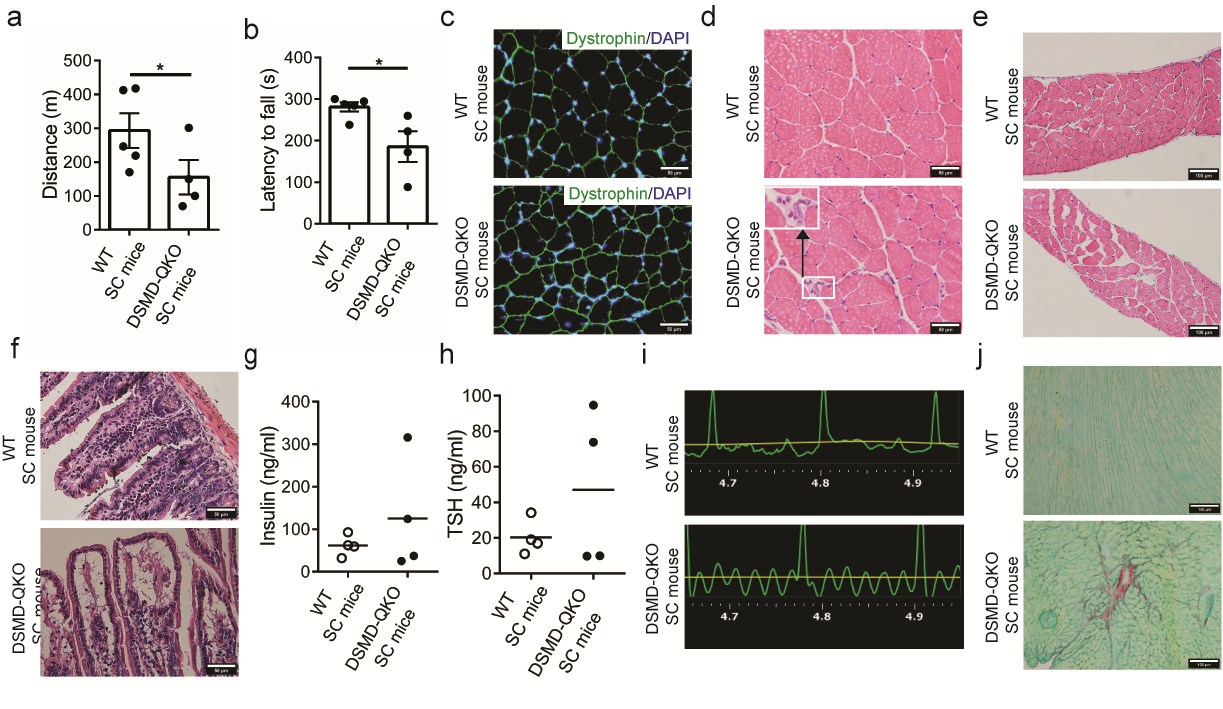

**Fig. S7** DM1-associated phenotypes in adult DSMD-QKO SC mice. **(a)** Treadmill test showed DSMD-QKO SC mice were weaker than WT SC mice (WT SC mice, n = 5; DSMD-QKO SC mice, n = 4). Unpaired Student’s *t* test, **P* < 0.05. **(b)** Rotarod analysis (WT SC mice, n = 5; DSMD-QKO SC mice, n = 4). Unpaired Student’s *t* test, **P* < 0.05. **(c)** Immunofluorescent staining indicated equivalent dystrophin levels in both DSMD-QKO and WT SC mice. **(d)** Representative images of H&E staining of TA muscle sections from DSMD-QKO SC mice and WT SC mice. White box indicated nuclear clumps in DSMD-QKO muscle. Scale bars, 50 μm. **(e)** Representative images of H&E staining of diaphragm showing loose construction and increased fat cell invasion in DSMD-QKO SC mice. Scale bars, 100 μm. **(f)**, H&E staining of small intestinal sections showing disorganization of intestine villi in adult DSMD-QKO mice. Scale bars, 50 μm. **(g-h)** Insulin and TSH levels were determined using ELISA showed increased hormone levels in some DSMD-QKO SC mice (n = 4 *per* group). **(i)** Electrocardiography experiments showed atrial flutter in DSMD-QKO SC mice. **(j)** Representative images of sirius red staining revealed myocardial fibrosis in DSMD-QKO SC mice. Scale bars, 100 μm.

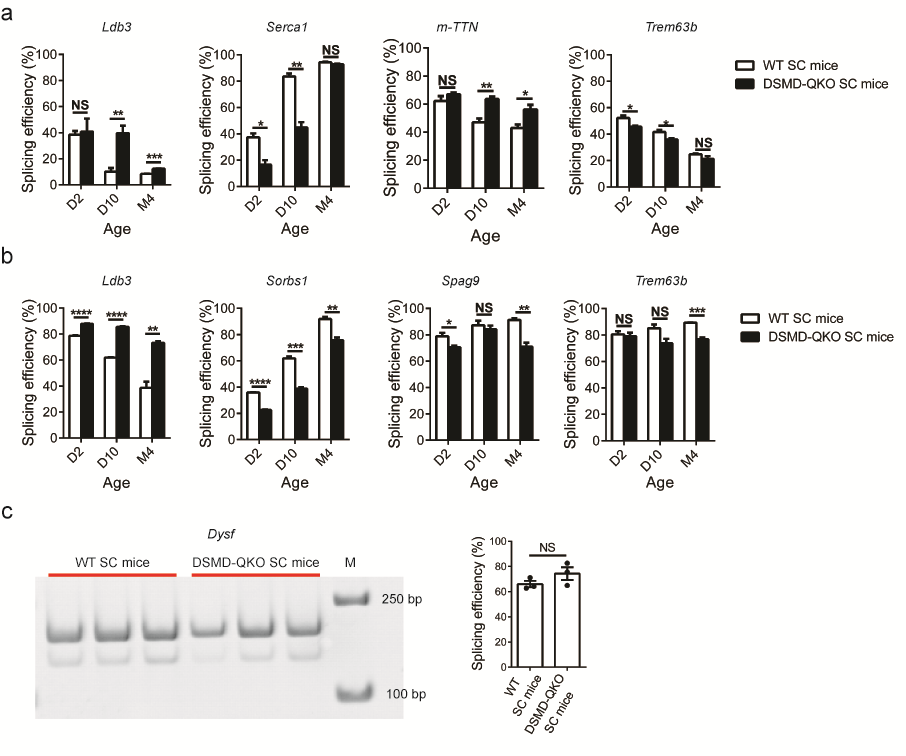

**Fig. S8** DM1-associated mis-splicing in adult DSMD-QKO SC mice. **(a)** Splicing efficiency for four target pre-mRNAs in TA muscles from DSMD-QKO SC mice and WT SC mice. Unpaired Student’s *t* test, **P* < 0.05, ***P* < 0.01, ****P* < 0.001. **(b)** Splicing efficiency for four target pre-mRNAs in cardiac muscle of DSMD-QKO SC mice and WT SC mice. Unpaired Student’s *t* test, **P* < 0.05, ***P* < 0.01, ****P* < 0.001, *****P* < 0.0001. **(c)** Splicing efficiency of *Dsyf* in TA muscles from DSMD-QKO SC mice and WT SC mice showed similar pattern, which also is not changed in muscles of human DM1 patients.

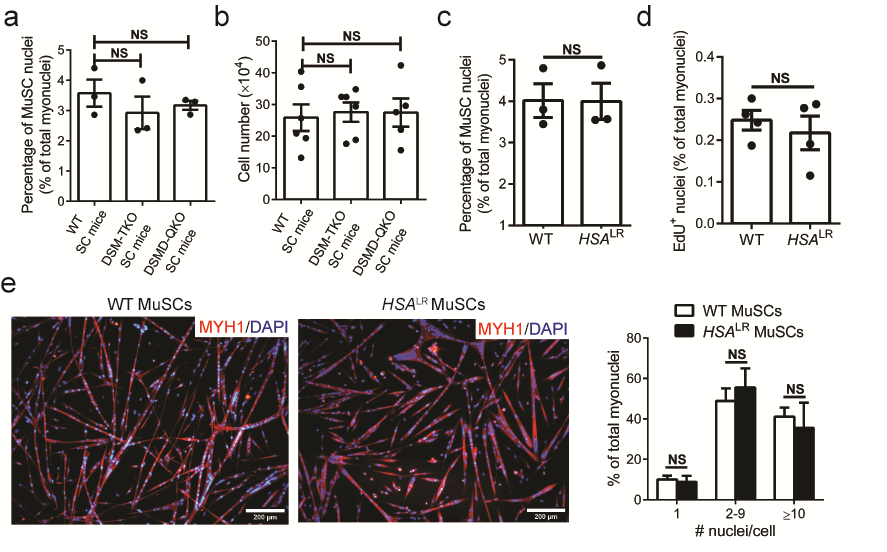

**Fig. S9** Differentiation defects in MuSCs of DSM-TKO and DSMD-QKO SC mice. **(a)** FACS analysis indicated equivalent number of satellite cells in DSM-TKO, DSMD-QKO and WT SC mice (n = 3 *per* group). Unpaired Student’s *t* test. NS, no significant differences. **(b)** The similar proliferation ability of cultured MuSCs from DSM-TKO, DSMD-QKO and WT SC mice (n = 3 *per* group). Unpaired Student’s *t* test. NS, no significant differences. **(c)** Equivalent number of satellite cells in *HSA^LR^* mice (FVB/n background) and WT mice (FVB/n background) (n = 3 *per* group). Unpaired Student’s *t* test. NS, no significant differences. **(d)** Equivalent in vitro proliferation ability of MuSCs isolated from *HSA^LR^* mice and WT mice (n = 3 *per* group). Unpaired Student’s *t* test. NS, no significant differences. **(e)** Equivalent in vitro differentiation potential of *HSA^LR^* and WT MuSCs. Scale bars, 200 µm. Unpaired Student’s *t* test. NS, no significant differences.

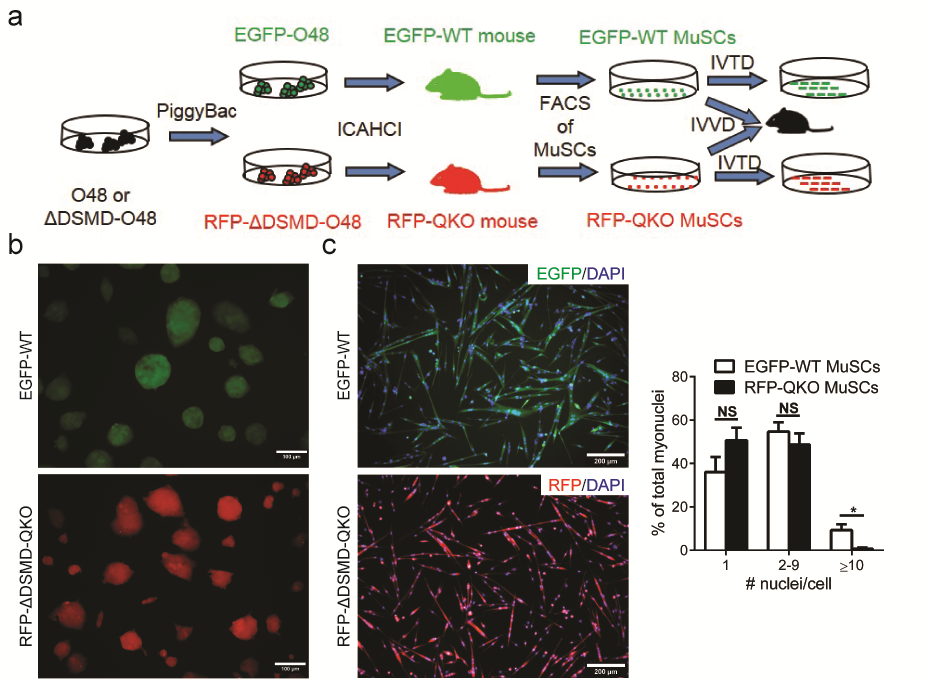

**Fig. S10** In vivo differentiation defects in MuSCs of DSMD-QKO SC mice. **(a)** Schematic diagram of isolation of DSMD-QKO MuSCs carrying RFP tansgenes (RFP-QKO MuSCs) and WT MuSCs carrying EGFP transgene (EGFP-WT MuSCs) generated by ICAHCI. IVTD, in vitro differentiation. IVVD, in vivo differentiation. **(b)** Transgenetic RFP-ΔDSMD-O48 and EGFP-O48 haploid ESCs, Scale bars, 100 µm. **(c)** In vitro differentiation of RFP-QKO MuSCs and EGFP-WT MuSCs. Unpaired Student’s *t* test. **P* < 0.05. NS, no significant changes. Scale bars, 200 µm.

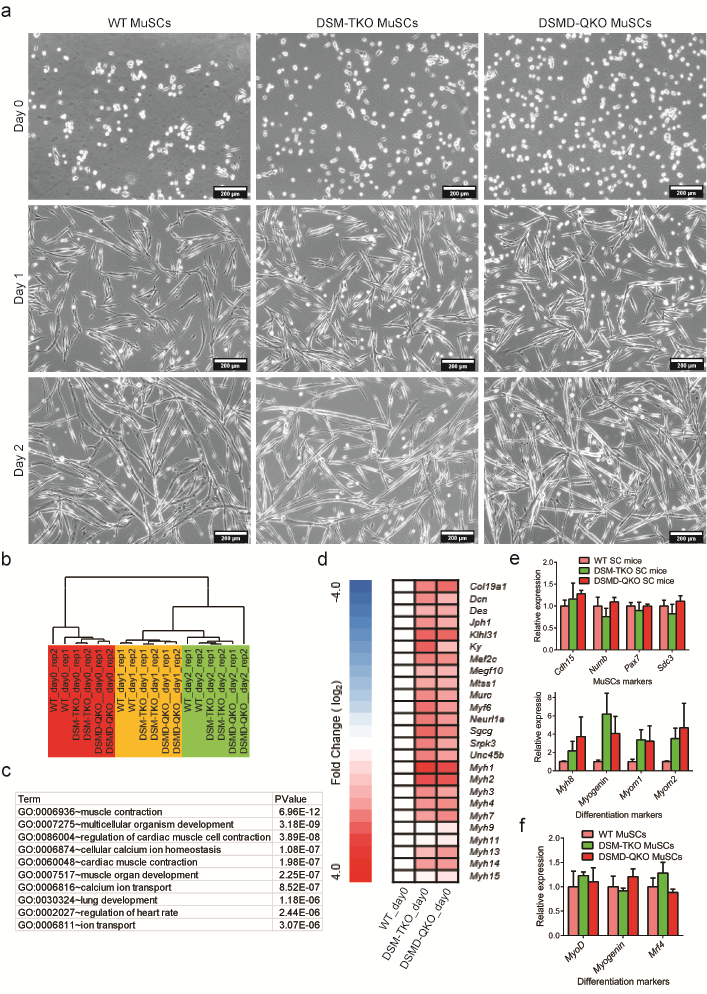

**Fig. S11** MuSCs of DSM-TKO and DSMD-QKO SC mice sustain less stemness. **(a)** Representative images of MuSCs (Day 0) and cells at the first and second day post differentiation of DSM-TKO, DSMD-QKO and WT SC MuSCs for RNA seqence. Scale bars, 200 µm. **(b)** Genomic expression profiles of MuSCs (Day 0) and cells at the first and second day post differentiation of DSM-TKO, DSMD-QKO and WT SC MuSCs. **(c)** GO analysis of upregulated genes in DSM-TKO and DSMD-QKO MuSCs compared with WT MuSCs. **(d)** Heat map analysis of differentiation markers between DM1 MuSCs and WT MuSCs. **(e)** RT-qPCR of representative MuSCs markers and differentiation markers in MuSCs. **(f)** RT-qPCR of representative differentiation markers in MuSCs differentiated for two days.

| **Haploid ESC lines used for ICAHCI (the number of cell lines tested)** | **Passage Number** | **No. of SC Embryos Transferred (2-cell)** | **No. of SC Pups (% of Transferred Embryos)** | **No. of SC pups** **surviving to P2 (% of birth)** | **No. of pups** **surviving to 3 weeks (% of P2)** | **No. of pups** **surviving to adult (% of P2)** |
| --- | --- | --- | --- | --- | --- | --- |
| O48 | P43-P51 | 430 | 55 (12.8) | 52 (94.5) | 52 (100) | 50 (96.2) |
| ΔDMPK-O48 (2) | P41-43 | 192 | 26 (13.5) | 24 (91.6) | 22 (91.6) | 21 (87.5) |
| ΔSIX5-O48 (2) | P40 | 156 | 15 (9.6) | 13 (86.7) | 13 (100) | 12 (92.3) |
| ΔMBNL1-O48 (2) | P40-P42 | 200 | 19 (9.5) | 19 (100) | 18 (94.7) | 18 (94.7) |
| ΔDMWD-O48 (2) | P46-47 | 330 | 31 (9.4) | 30 (96.8) | 26 (86.7) | 25 (83.3) |
| DSM-O48 (2) | P57-P62 | 270 | 34 (12.6) | 32 (94.1) | 30 (93.8) | 30 (93.8) |
| DSMD-O48 (3) | P66-71 | 372 | 45 (12.1) | 35 (77.8) | 19 (54.3) | 17 (48.6) |

Table S1. Development of ICAHCI embryos derived from haploid ESCs with single, triple or quadruple mutations.

| **sgRNAs of targeted genes** | **No. of mismatch** | **No. of found targets (gene)** | **No. of mutated sites** | **Cell lines tested** |
| --- | --- | --- | --- | --- |
| Dmpk | 0  1  2  3  4 | 1 (*Dmpk*)  0  1  21  275 | 1  0  0  0  0 | △DSM-O48-1  △DSMD-O48-2 |
| Six5 | 0  1  2  3  4 | 1 (*Six5*)  0  0  16  186 | 1  0  0  0  0 | △DSM-O48-1  △DSMD-O48-2 |
| Mbnl1-up | 0  1  2  3  4 | 1 (*Mbnl1*)  0  1  11  220 | 1  0  0  0  0 | △DSM-O48-1  △DSMD-O48-2 |
| Mbnl1-down | 0  1  2  3  4 | 1 (*Mbnl1*)  0  0  13  198 | 1  0  0  0  0 | △DSM-O48-1  △DSMD-O48-2 |
| Dmwd | 0  1  2  3  4 | 1 (*Dmwd*)  0  4  40  415 | 1  0  0  0  0 | △DSM-O48-1  △DSMD-O48-2 |

**Table S2 off target analysis.**

|  | **Skeletal muscle** | | | | **Cardiac problems** | **Cataract** | **Endocrine disorders** | **Digestive system dysfunction** | **Congenital DM1** | | | **Defective satellite cells** | **References** |
| --- | --- | --- | --- | --- | --- | --- | --- | --- | --- | --- | --- | --- | --- |
|  | **Myotonic** | **Wasting** | **Weakness** | **Histology** |  |  |  |  | **Hypotonia** | **Breathing problem** | **Developmental delay** |  |  |
| Clinical features | +++ | +++ | ++ | +++ | ++ | ++ | ++ | ++ | ++ | ++ | ++ | + | (*11*) |
| *DMPK*^-/-^ | - | - | - | + | - | - | ND | ND | - | - | - | ND | (*12**,* *13*) |
| *SIX5*^-/-^ | - | - | - | - | - | + | ND | ND | - | - | - | ND | (*14**,* *15*) |
| *HSA*^LR^ (TG) | +++ | - | - | ++ | - | - | ND | ND | - | - | - | ND | (*16*) |
| *MBNL1*^-/-^ | +++ | - | - | ++ | - | +++ | ND | ND | - | - | - | ND | (*17*) |
| *MBNL1*^-/-^; *MBNL2*^C/C^; *Myog-Cre*^+/-^ | ND | ++ | +++ | +++ | ND | ND | ND | ND | - | + | + | ND | (*18*) |
| *MBNL1*^-/-^; *MBNL2*^C/C^; *MBNL3* ^C/Y^; *Myog-Cre*^+/-^ | ND | +++ | +++ | +++ | ND | ND | ND | ND | + | ++ | ++ | ND | (*18*) |
| *DMPK*^+/-^ | - | - | ND | - | - | - | - | - | ND | - | - | - | In this study |
| *SIX5*^+/-^ | ND | - | ND | - | - | - | - | + | ND | - | - | - | In this study |
| *MBNL1*^+/-^ | ND | - | ND | - | - | - | - | - | ND | - | - | - | In this study |
| *DMWD*^+/-^ | ND | + | ND | - | - | - | - | - | ND | - | - | - | In this study |
| DSM-TKO | + | ++ | ++ | + | + | - | + | + | - | - | - | + | In this study |
| DSMD-QKO | + | ++ | ++ | + | ++ | + | + | + | ++ | ++ | + | + | In this study |

Table S3. Clinical features of DM1 and phenotypes in mouse disease models.

-: No phenotype; +: Mild phenotype; ++: Moderate phenotype; +++: Severe phenotype.; ND: No determined.
